# Characterization of CAR T Cells Manufactured using Genetically Engineered Artificial Antigen Presenting Cells

**DOI:** 10.1101/2023.06.28.546908

**Authors:** Ali Sayadmanesh, Mohammad Azadbakht, Kheirollah Yari, Ali Abedelahi, Hajar Shafaei, Dariush Shanehbandi, Behzad Baradaran, Mohsen Basiri

## Abstract

**Objective:** Chimeric antigen receptor (CAR) T cell therapy has recently emerged as a promising approach for the treatment of different types of cancer. Improving CAR T cell manufacturing in terms of costs and product quality is an important concern for expanding the accessibility of this therapy. One proposed strategy for improving T cell expansion is to use genetically engineered artificial antigen presenting cells (aAPC) expressing a membrane-bound anti-CD3 for T cell activation. In this study, we characterized CAR T cells generated with this approach in terms of expansion efficiency, immunophenotype, and cytotoxicity.

**Materials and Methods:** In this experimental study, we generated an aAPC line by engineering K562 cells to express a membrane-bound anti-CD3 (mOKT3). T cell activation was performed by culturing PBMCs with either mitomycin C-treated aAPCs or surface-immobilized anti-CD3 and anti-CD28 antibodies. Untransduced and CD19-CAR-transduced T cells were characterized in terms of expansion, activation markers, IFN-γ secretion, CD4/CD8 ratio, memory phenotype, and exhaustion markers. Cytotoxicity of CD19-CAR T cells generated by aAPCs and antibodies was also investigated using a bioluminescence-based co-culture assay.

**Results:** Our findings showed that the engineered aAPC line has the potential to expand CAR T cells similar to that of the antibody-based method. Although activation with aAPCs leads to a higher ratio of CD8^+^ and effector memory T cells in the final product, we did not observe a significant difference in IFN-γ secretion cytotoxic activity or exhaustion between CAR T cells generated with aAPC or antibodies.

**Conclusion:** Our results show that despite the differences in the immunophenotypes of aAPC and antibody-based CAR T cells, both methods can be used to manufacture potent CAR T cells. These findings can be instrumental for the improvement of the T cell manufacturing process and future applications of aAPC-derived CAR T cells.

## Introduction

Adoptive Cell therapy (ACT) is a type of personalized immunotherapy in which immune cells are isolated and expanded outside the body, or sometimes modified, and then returned to the patient to boost the body’s fight against diseases such as cancer (1–3). Chimeric antigen receptor (CAR) T cell therapy, as a type of adoptive cell transfer, is an innovative and promising approach in the field of cancer immunotherapy (4–6). CAR T cells are autologous T cells engineered to express target antigen-specific receptors with an intracellular CD3 domain fused to a stimulatory domain that does not require antigen-presenting cells to target malignant cells (7, 8). CAR T cells targeting CD19 and B-cell maturation antigen (BCMA) are currently being used to treat some hematological malignancies, including acute lymphoblastic leukemia (B-ALL), large B-cell lymphoma (LBCL), Follicular lymphoma (FL), multiple myeloma (MM) and mantle cell lymphoma (MCL) have been approved in the United States and Europe(6, 9, 10).

*Ex vivo* expansion of T cells requires activation through TCR and co-stimulatory signals which is conventionally performed by bead- or surface-immobilized anti-CD3 and anti-CD28 antibodies. However, challenges such as the cost of manufacturing CAR T cells, disposable reagents with GMP quality, long-term growth of pure antigen-specific and polyclonal CD8^+^ T cells, and immunological synapse formation are the main obstacles to the clinical application of CAR T therapies. Therefore, one of the strategies aimed at improving the expansion and activation of T cells and minimizing the existing challenges has been the use of aAPCs to produce engineered T cells for clinical applications (11–13). One of the common approaches in this field is the use of the K562 cell line. Unlike anti-CD3/CD28 antibodies, the K562 aAPC approach is cost-effective and may include additional stimulatory molecules such as ICAM-1 and LFA-3 that are more capable of stimulating and interacting with T cells (14, 15). Other advantages of K562 aAPCs include the lack of MHC expression, the ability to grow in a serum-free environment, expand the CD8^+^ T cell population from donors in a short period of time, and the possibility of better immunological synapse formation due to the fluidity of the cell membrane (16, 17). Also, in previous studies conducted on humans, the use of K562 cell has been investigated in terms of safety (18, 19). However, characterization and comparison of aAPC- and antibody-expanded T cells in terms of immunophenotype and functional properties is crucial to understand potential differences between these two methods.

In this study, we engineered a K562 aAPC that can be used to expand, transduce, and generate CAR T cells without the need of anti-CD3/CD28. Our K562 aAPC expressing a membrane-bound anti-human CD3ε (OKT3) single-chain variable fragment (scFv) efficiently expanded T cells suitable for viral transduction. We confirmed that our aAPC significantly expanded CD8^+^ T cells compared to the Ab activation method, and CAR T cells generated with this aAPC had a similar effect on T cell exhaustion markers as the standard method. Finally, we demonstrated that this method of producing CAR T cells is comparable to the standard method in terms of in vitro antitumor activity.

## Materials and Methods

### Constructs preparation

A DNA sequence coding for a fusion protein consisting of of human CD5 signal sequence(nucleotines 73 to 135 of the GenBank accession number NM_014207), OKT3 scFv (nucleotines 64 to 810 of the GenBank accession number HM208751), and human CD8A hinge and transmembrane regions (nucleotines 1301 to 1516 of the GenBank accession number NM_001768) was synthesized (ShineGene, China) and recived in pUC57 plasmid. TThis fragment then was subcloned into a retroviral SFG.CNb30_opt.IRES.eGFP (Addgene, USA) at the NcoI and XhoI restriction sites. The CD19-CAR sequence was then cloned into SFG.CNb30_opt.IRES.eGFP at the NcoI and Kpn21 restriction sites. The CD19-CAR construct consisted of CD19-specific FMC63 mAb single-chain variable fragment, a CD8 hinge region, CD28 transmembrane, and cytoplasmic regions, and the cytoplasmic region of CD3 zeta in a retroviral backbone was obtained from Royan Institute Plasmid Bank.

### Cell lines

K562 and Raji cell lines were obtained from the American Type Culture Collection (ATCC, Manassas, VA, USA). K562 and Raji cells were maintained in RPMI 1640 and supplemented with 10% FBS, GlutaMax (1X), penicillin (100 units/ml), and streptomycin (100 μg/ml) (Gibco) at 37°C and 5% Co2. The Platinum-Amphotropic (Plat-A) retroviral packaging cell line (Cell Biolabs, San Diego, CA, USA) was maintained in DMEM High Glucose (Dulbecco’s Modified Eagle Medium) and supplemented with 10% FBS, GlutaMax (1X), penicillin (100 units/ml) and streptomycin (100 μg/ml, Gibco).

### Retrovirus production

Retroviral particles were separately produced by transient transfection of Plat-A cells with the OKT3 or CD19-CAR-encoding SFG plasmids (18μg) in 100 mm tissue culture plates using Lipofectamine™ 3000 reagent (L3000075, Invitrogen, USA). Media were refreshed 16 hours post-transfection and supernatants containing retroviral particles were collected 48-72 hours after transfection. Viral supernatants were filtered through a 0.45 μm filter and used for transduction experiments.

### aAPC line (OKT3-K562 cell line) production

K562 cells were cultured in non-tissue culture 24-well plate at the density of 2 × 10^5^ per well. After an overnight cells were centrifuged in 400g and re-suspended in OKT3-retroviral in the presence of 8μg/ml Polybrene (Santa Cruz Biotechnology, sc-134220). After 12 hours, the medium was replaced with a fresh complete medium and incubated for an additional 2 days at 37°C with 5 % CO2, and GFP expression was determined by fluorescent microscope (Olympus, IX71). Transduced cells were cultured for 10 days (five passages) and then subjected to fluorescence-activated cell sorting (FACS) to obtain a uniform population of GFP-positive cell designed as OKT3-K562 cells (aAPC line).

### Treatment aAPC with mitomycin C

The aAPCs were seeded in non-tissue culture 24-well plate at the number of 1 × 10^6^ cells per well in a volume of 1 ml complete medium and were treated with 20μg mitomycin C at different times (30, 60 and 90 minutes). In order to investigate the effect of mitomycin C on cell proliferation, cells were washed with PBS and re-suspended in a complete medium, and counted for 4 consecutive days.

### Co-culture of aAPCs with PBMCs

Initially, human PBMCs (peripheral blood mononuclear cells) were isolated from healthy donors using Ficoll-Paque density gradient centrifugation (Progen, 1114544). In each well of a 24-well plate, 1 × 10^6^ isolated PBMCs were co-cultured with 1 × 10^6^, 5 × 10^5^, or 2.5 × 10^5^ mitomycin C-treated aAPCs, corresponding for the PBMC:aAPC ratios of 1:1, 1:0.5, and 1:0.25 respectively. On day 1, CTL medium containing 100 IU/ml human IL-2 (R&D Systems, 202-IL-010/CF) was added to the plates and the expansion of T cells continued. Every two days, the cells were counted with a Neobar slide and splitted. Also, to determine whether T cell killed mitomycin C-treated aAPCs, they were co-cultured with PBMC cells at the ratio of 1:1, 1:0.5, and 1:0.25 (PBMC:aAPC). Considering that aAPCs express the GFP marker, on days 0, 1, and 2 the killed aAPCs were analyzed by flow cytometry. As a control group, 1 × 10^6^ PBMCs cells were cultured in a 24-well plate pre-coated with anti-CD3 (Miltenybiotec, 130-093-387) and anti-CD28 (Miltenybiotec, 130-093-375) antibodies.

### CAR T cell generation

To stimulate T cells, PBMCs were first isolated and divided into two groups, one group was cultured in a non-tissue culture 24-well plate pre-coated with anti-CD3 and anti-CD28 antibodies, and the other group was co-cultured with mitomycin C-treated aAPCs at the ratio of 1:0.5 (1 × 10^6^ PBMCs: 0.5 × 10^6^ aAPCs) in 2 ml/well CTL medium containing RPMI1620 (Cytiva, SH3009601) and Click’s medium (Fujifilm Irvine Scientific, 9195) in equal proportion supplemented with 1% GlutaMax (Gibco, 35050061) and 10% FBS (Gibco, 16140071). On day 1, aspirated 1 ml of media and added 1 ml of CTL media containing 200 IU/ml human IL-2 (R&D Systems) to a final concentration of 100 IU/ml. On day 3, the non-tissue culture 24-well plates were coated with 7 μg/ml RetroNectin (TAKARA) for 3 hours at 37°C. Aspirated RetroNectin solution and 2 ml anti-CD19-CAR retroviral supernatants were added to each well and centrifuged at 2000g at 4°C for 90 minutes. Activated T cells were re-suspended at the concentration of 2 × 10^5^ cells/well in 2 ml CTL media, in the final concentration of 100 IU/ml human IL-2, and centrifuged at 400g for 10 minutes. During T cell expansion, cells were split every two days with a CTL medium containing a final concentration of 50 IU/mL of IL-2.

### Flow cytometry

T cells were staining 30 min at 4°C with anti-human CD3-PE (Cyto Matin Gene, CMGCD3-P100) anti-CD3-PerCP (BD, 347344), anti-CD3-FITC (BD, 349201), anti-CD4-FITC (BD, 555346), anti-CD8-PE (Cyto Matin Gene, CMGCD8-P), anti-CD25-FITC (BioLegend, 302603), anti-CD69-PE (BioLegend, 310906), Fc-CD19-FITC (abcam, ab246020), anti-TIM-3-PE (BD, 583422), anti-PD-1-FITC (BioLegend, 329903), anti-CCR7 (BioLegend, 353216), anti-CD45RA (BD, 556627), CD56-PE (BD, 555516). Samples were washed with PBS and flow cytometry acquisition was performed on BD FACSCalibur (BD Biosciences, USA). 7-AAD was used for staining the dead cells (eBioscience, 00-6993-50). Analysis was performed using FlowJo software (Treestar, USA).

### In vitro cytotoxicity assay

Cytotoxic activity of CAR T cells against target cells was determined by a luciferase-based cytotoxicity assay and flow cytometry-based cytotoxicity assay. The CAR T cells were incubated with ffluc-expressing Raji cells at various E:T ratios in a flat bottom 96-well plate. After 6 h, added 100 μl of 75 μg/ml of sigma D-luciferin (L6882, Sigma) to each well, and luciferase activity was measured immediately with an Alliance Q9 Advanced imaging system (UVITEC, UK). Untransduced T cells with ffluc-expressing Raji cells and ffluc-expressing Raji cells without effector cells served as controls. The percentage of the specific lysis was calculated according to the following formula: % specific lysis = 100 × (spontaneous cell death RLU-sample RLU)/(spontaneous death RLU-maximal killing RLU). For flow cytometry-based T cell cytotoxicity assay, CAR T cells, and untransduced T cells were added at a 1:5 effector to tumor cell ratio (25 × 10^4^ T cells: 125 × 10^5^ Raji cells) in not-treated 6-well (Corning, 3736) and wells without effector cells served as untreated controls. Cells were co-cultured for 6 days in the presence of 2 mL complete media and on days 0, 3, and 6, and cells were harvested and stained with anti-CD3-PE. As previously described (20) for quantifying cells by flow cytometry, 20 μL of CountBright™ Absolute Counting Beads (C36950; Invitrogen, Eugene, OR) were added, and for excluding the dead cells, 7-AAD was added. The acquisition was halted at 2000 beads.

### Assessment of activation markers expression and IFN-γ release assay

CAR T cells and untransduced T cells prepared with aAPC and anti-CD3/CD28 were co-cultured with Raji cells at a ratio of 1:1 (1 × 10^5^ cells) in a 24-well plate. After 24 hours, supernatants were harvested to measure the release of IFN-γ using a IFN-γ-specific ELISA according to the manufacturer’s instructions (R&D Systems, DY28B05). To evaluate T cell activation markers, cells were stained with anti-CD25 and anti-CD69 antibodies and were measured by flow cytometry.

### Statistical analyses

Statistical analysis was performed using GraphPad Prism software (V.8.4.1) and P-values were calculated by ordinary one-way ANOVA Tukey’s multiple comparisons test for Fig. 3B and Two-way ANOVA with Tukey’s multiple comparisons test for 3C, 4F, 5B, 6B, 6E, and 6F. For 3D, 3E, 3F, 4G, 4H, 4I, and 5D P-values were calculated with two-way ANOVA with Sidak’s multiple comparisons test and for 4D with ordinary one-way ANOVA with Sidak’s multiple comparisons test. In addition, Fig. 3G and 4J used an unpaired t-test. P-values less than 0.03 was considered statistically significant and is marked as * in the figure legends. All results are represented as mean ± standard deviation (SD).

## Results

### Generation of the aAPC line

To generate an aAPC line expressing mOKT3, we transduced K562 cells with a retroviral vector encoding mOKT3 together with a GFP reporter (Fig. 1A). To confirm the transduction, K562 cells were examined for GFP expression using a fluorescent microscope 48 hours after transduction. After five passages, GFP-positive K562 cells were sorted by FACS to establish the aAPC line. Afterward, GFP expression was confirmed by fluorescent microscopy and flow cytometry in the aAPC line (sorted K562 transduced cells) in comparison with unsorted K562 transduced cells and untransduced K562 cells (Fig. 1B). Results showed that the unsorted cells were transduced with an efficiency of about 10% and after sorting, 93% of them were positive for the GFP reporter.

**Figure 1.**
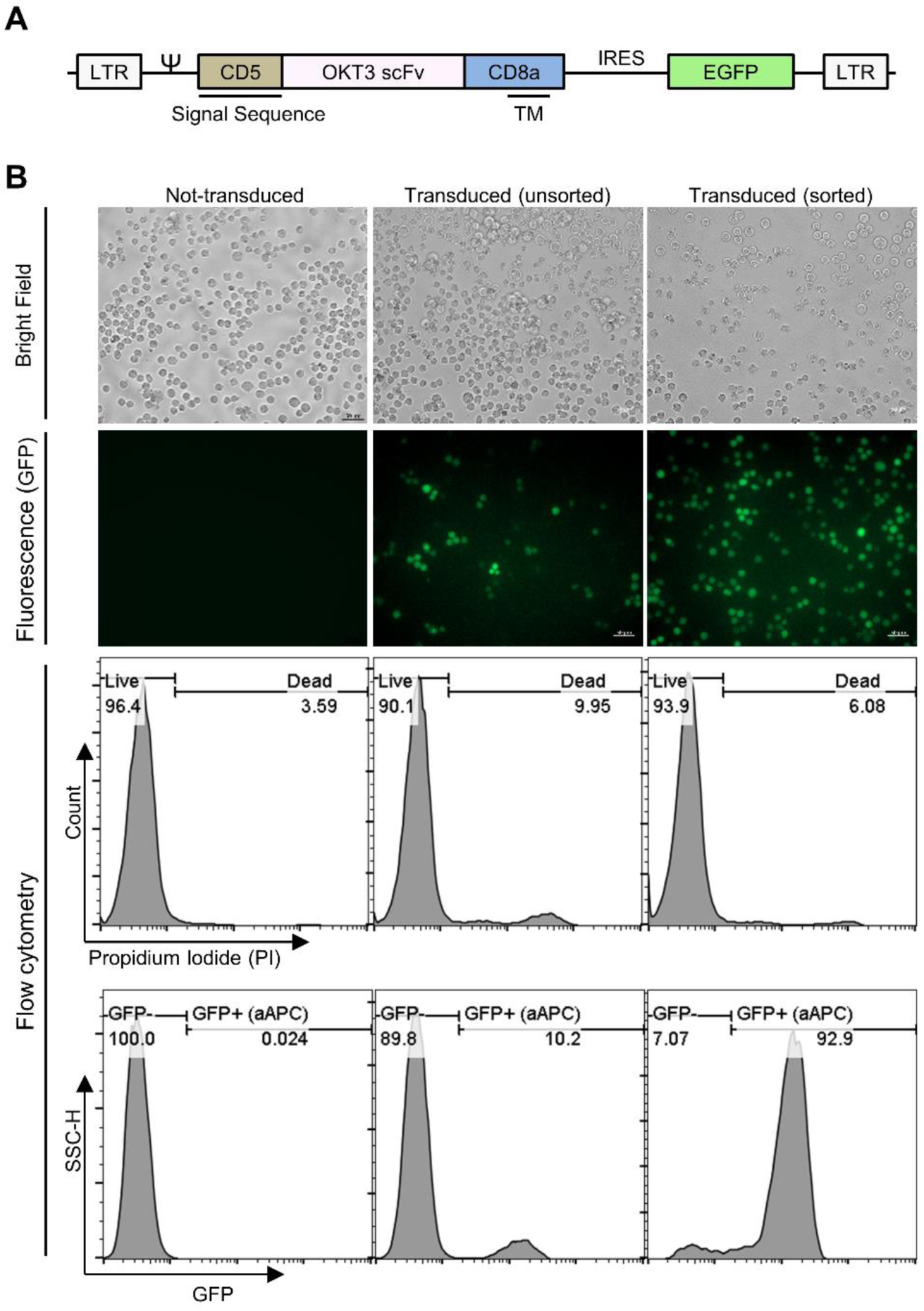
mOKT3 construct and generation of aAPC cell line. (A) Schematic presentation of the mOKT3 construct containing a CD5 signal sequence, a synthetic gene coding for the OKT3 scFv, a CD8a transmembrane region, and IRES-EGFP in a SFG retroviral vector. (B) OKT3 transduction on the K562 cell line was confirmed. The first and second panels from the top respectively: Bright field and GFP fluorescent images of the non-transduced, transduced, and sorted OKT3-K562 cell line by fluorescent microscopy. Third panel: Total cell viability of corresponding cells before GFP+ gating. Fourth panel (lower panel): Transduction efficiency was confirmed by flow cytometry.

### Co-culture of aAPC with primary PBMC

In order to use the aAPCs for T cell activation, we treated them with mitomycin C in different time point and cultured them for 4 days. The results of cell counting did not show any difference between the times of cell treatment with mitomycin C, so the shortest time (30 minutes) was chosen for the incubation time in the rest of the experiments (Fig. 2A). To investigate cluster formation of aAPC and T cells we co-cultured PBMCs and mitomycin C-treated aAPCs in the ratio of 1:1, 1:0.5, and 1:0.25 (PBMC:aAPC) and one group activated with anti-CD3/CD28 as a standard T cell activation protocol (Fig. 2B). The results showed that at higher aAPC ratios, more clusters were formed and the 1:0.5 ratio was similar to the standard T cell activation method in terms of the amount of cluster formation (Fig. 2C). Therefore, it seems that this ratio is suitable for cell activation. To measure the survival time of aAPCs in contact with T cells, we measured the survival of aAPCs at the respective ratios on days 0, 1, and 2 (Fig. 2D). Results showed on day 2 after co-culture in the 1:1 ratio more aAPCs are still alive, but at 1:0.5 and 1:0.25 ratios few aAPCs remain (Fig. 2E and F). Overall, according to the obtained results, the ratio of 1:0.5 was the most appropriate in terms of the cluster formation, and also considering that T cell transduction is performed on day 3 in most CAR T cell production protocols, the aAPCs did not persist in the culture after 3 days. Therefore, in the rest of experiments, the ratio of 1:0.5 was chosen as a suitable ratio for the activation of T cells.

**Figure 2.**
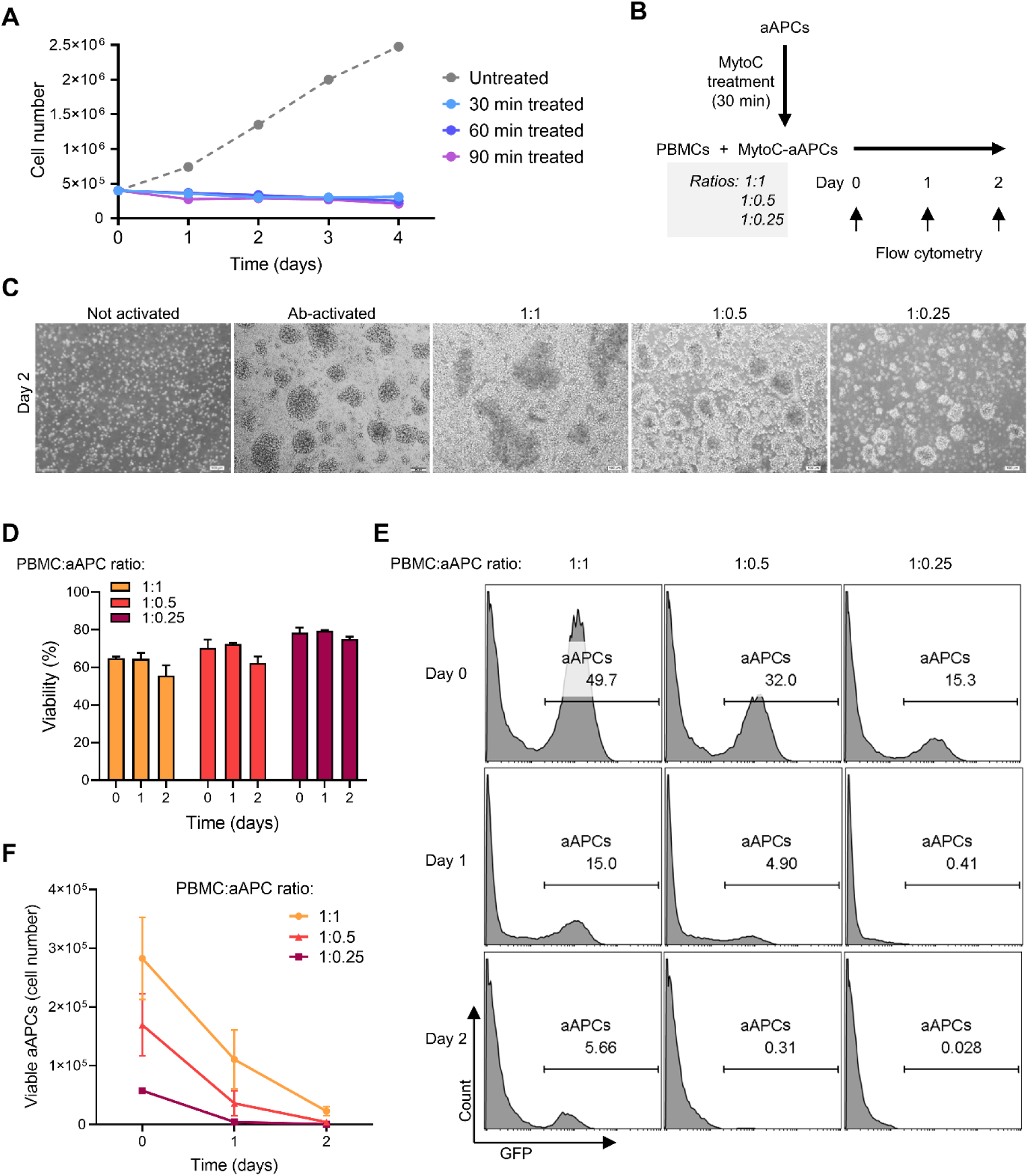
Co-culture of aAPC cells with PBMCs and their viability assay. (A) Treatment of aAPC with mitomycin C (MytoC) for different durations showed that treated aAPC cell lines did not show proliferation. The data showed that aAPC cell lines treated with mitomycin C did not show proliferation for up to 4 days. *n* = 1 (B) Schematic of PBMC and aAPC coculture in different ratios of 1:1, 1:0.5, and 1:0.25. (C) Cluster formation of T cells after PBMCs co-cultured with mitomycin C treated-aAPC cells in different ratios (1:1, 1:0.5, and 1:0.25) and compared with standard T cell activation with anti-CD3/antiCD28 antibodies (Ab). (D) Viability of cocultured cells after on day 2. (E) Representitive data showing GFP expression among the viable cells in cocultures, where GFP^+^ cells represent aAPCs. The co-culture of aAPC cells with PBMCs at 3-time points showed that, on day 2, aAPC cells were eliminated in the ratio of 1:0.5 and 1:0.25, but in the ratio of 1:1, about 5% of the aAPC cells remained in the culture. (F) Summery of data showing the quantified viable aAPC cells in different coculture conditions. The ratios of 1:0.5 and 1:0.25 on day 2, the number of aAPC cells was greatly reduced and almost the majority of cells in the culture were eliminated. *n* = 3.

### T cell expansion after aAPC-mediated activation

In order to evaluate the rate of T cell expansion with aAPC, we cultured PBMCs activated with aAPC, K562 (negative control), or anti-CD3/CD28 (positive control) for seven days in the presence of IL-2 (Fig. 3A) The cell counting results showed that PBMCs activated with aAPCs could expand similarly to the Ab activated method (anti-CD3/CD-28 solution) and increased about 22-fold within 7 days. Also, lower rates of cell expansion were observed with K562 cells (Fig. 3 B). To determine the cell types expanded in the experimental groups, we used the CD3 and CD56 markers. Our data showed that the aAPC activation resulted in a predominant CD3^+^ T cell population similar to the anti-CD3/CD-28 group (Fig. 3C). The results also indicated that most of the cells expanded after K562 co-culture are CD3^−^CD56^+^ NK cells (Fig. 3C).

**Figure 3.**
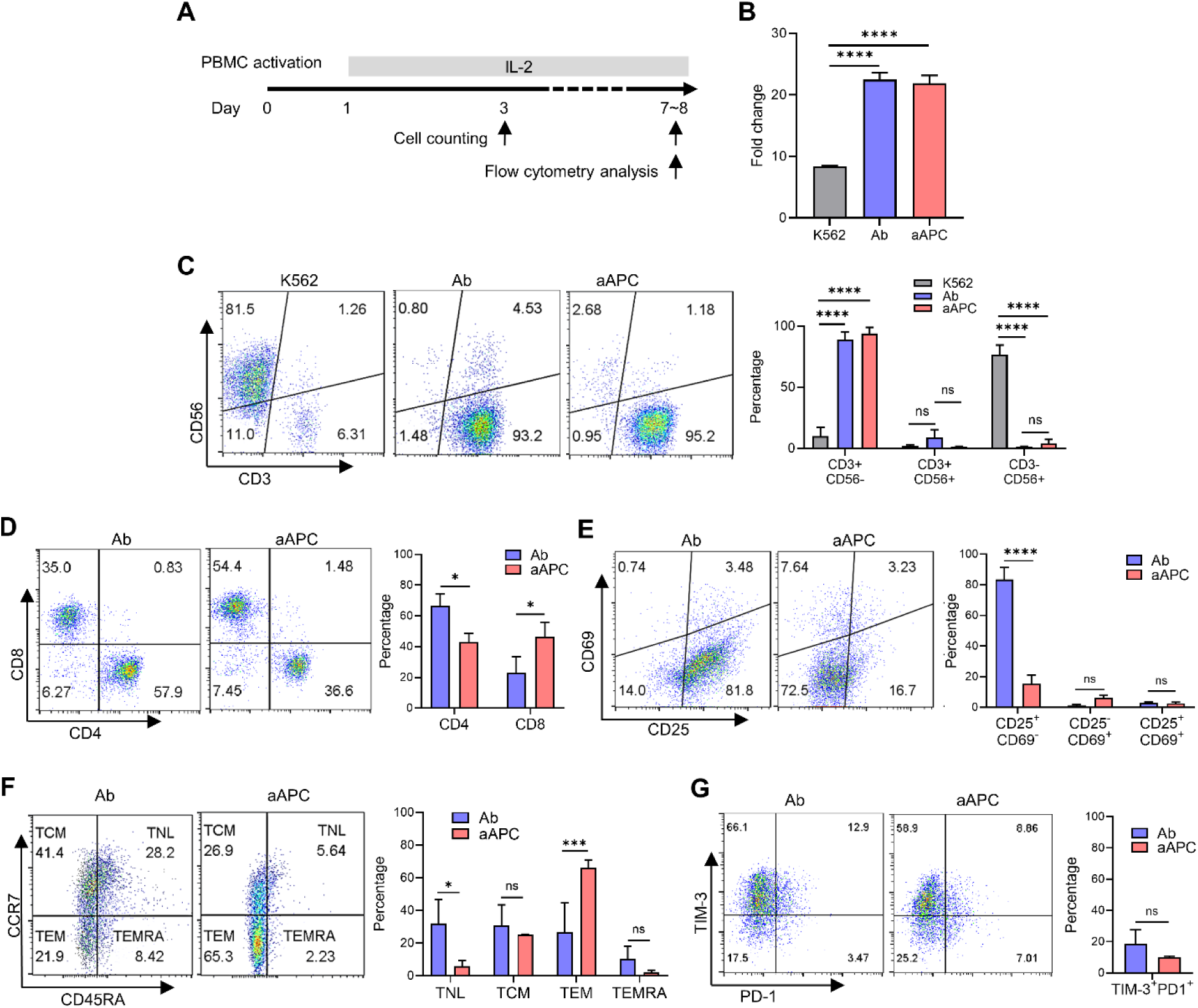
Expansion of T and NK cells and evaluation of markers of activation, exhaustion, and immunophenotype of T cells in aAPC activation method. (A) Schematic illustration of the experiment for T cell activation with aAPC cells, anti-CD3/CD28 antibodies, and K562 cell line and cell expansion and staining. (B) Fold expansion of activated T cells with aAPC cells, anti-CD3/CD28 antibodies (Ab), and K562 cell line as a negative control. Cell expansion was assessed from day 3 to day 7 and observed a 22-fold expansion at 4 days. (C) Cells were stained with anti-CD56 and anti-CD3 to determine of T cell and NK cell populations. The aAPC cell could expand T cells (CD3^+^CD56^−^) similarly to the Ab activated method and in the two groups with no significant variation in the expansion of the NKT population (CD3^+^ CD56^+^). K562 cell line was not suitable for T cell expansion, but it was able to expand NK cells (CD3^−^CD56^+^). (D) Analysis of T cells to determine the CD4/CD8 ratio showed that CD8^+^ T cells were more expanded in the aAPC activated method and CD4^+^ T cells in the Ab activated method. (E) To assess CD25 and CD69 activation markers, cells were stained and measured by flow cytometry. The data showed no difference between the expression of CD25^+^CD69^+^ dual activation markers in aAPC and Ab activated method and the expression of CD25^+^CD69^−^ marker increased in Ab activated method. (F) Representative flow cytometry and quantification data for memory phenotype analysis of T cells on day 8. The aAPC activated method increased the differentiation of T cells into TEM cells, and the Ab activated method increased the differentiation of T cells into TNL cells. (G) The expression of PD-1 and TIM-3 exhaustion markers in different methods of activation measured by flow cytometry showed that T cell exhaustion markers were not significant in both groups. TNL: naïve-like T cell; TCM: central memory T cell; TEM: effector memory T cell; TEMRA: CD45RA^+^ effector memory T cell. *n* = 3. * P < 0.05; *** P < 0.001; **** P < 0.0001.

### Immunophenotype of the T cells expanded with aAPC

To evaluate of effect of different activation methods on T cell immunophenotype, we stimulate PBMCs cells with aAPCs or anti-CD3/CD28 similar to the previous experiment (Fig. 3A) and assessed the expression of different T cell markers after 7-8 days. We analyzed CD4 and CD8 markers to assess the effect of aAPC and Ab activation methods on T cell subpopulations. The analysis revealed that in the aAPC-activated group, CD8^+^ T cytotoxic cells expanded more than CD4^+^ T helper, instead, in the Ab-activated group, CD4^+^ T helper cells expanded more than CD8^+^ T cytotoxic cells (Fig. 3D). Investigation of the activation phenotype of T cells with CD25 and CD69 staining showed no difference in the CD25^+^CD69^+^ and CD25^−^CD69^−^ population between aAPC and Ab groups, while the proportion of CD25^+^CD69^−^ population was decreased aAPC compared with Ab group (Fig. 3E). To compare the role of various activation methods on memory phenotype of T cells, the expression of CD45RA and CCR7 were measured with flow cytometry. As the result shows in Fig. 3F, aAPC-mediated activation increased the differentiation of T cells to T effector memory cells (TEM; CD45RA^−^CCR7^−^) compared with Ab-mediated activation. Correspondingly, the proportion of naïve-like (TNL; CD45RA^+^CCR7^+^) sub-population was higher in the Ab-activated T cells. However, the percentages of central memory T cells (TCM; CD45RA^−^CCR7^+^) and CD45RA^+^ effector memory T cells (TEMRA; CD45RA^+^CCR7^−^) were not significantly different between Ab and aAPC groups (Fig. 3F). We did not observe any statistically significant difference in the expression of exhaustion markers PD-1 and TIM-3 between T cells activated by Ab and aAPC (Fig. 3G).

### CAR T cell generation by aAPC-mediated activation

In order to evaluate the application of aAPC line for manufacturing clinically relevant T cell products, we used a retroviral second-generation CD19-specific CAR construct to generate CD19 CAR T cells (Fig. 4A and B). We were able to achieve transduction efficiencies higher than 80% with both aAPC-activated and Ab-activated T cells (Fig 4. C). The expansion of the CAR transduced T cells in both aAPC and Ab groups where significantly higher than that of T cells cultured with K562 which served as a negative control (Fig. 4D).

**Figure 4.**
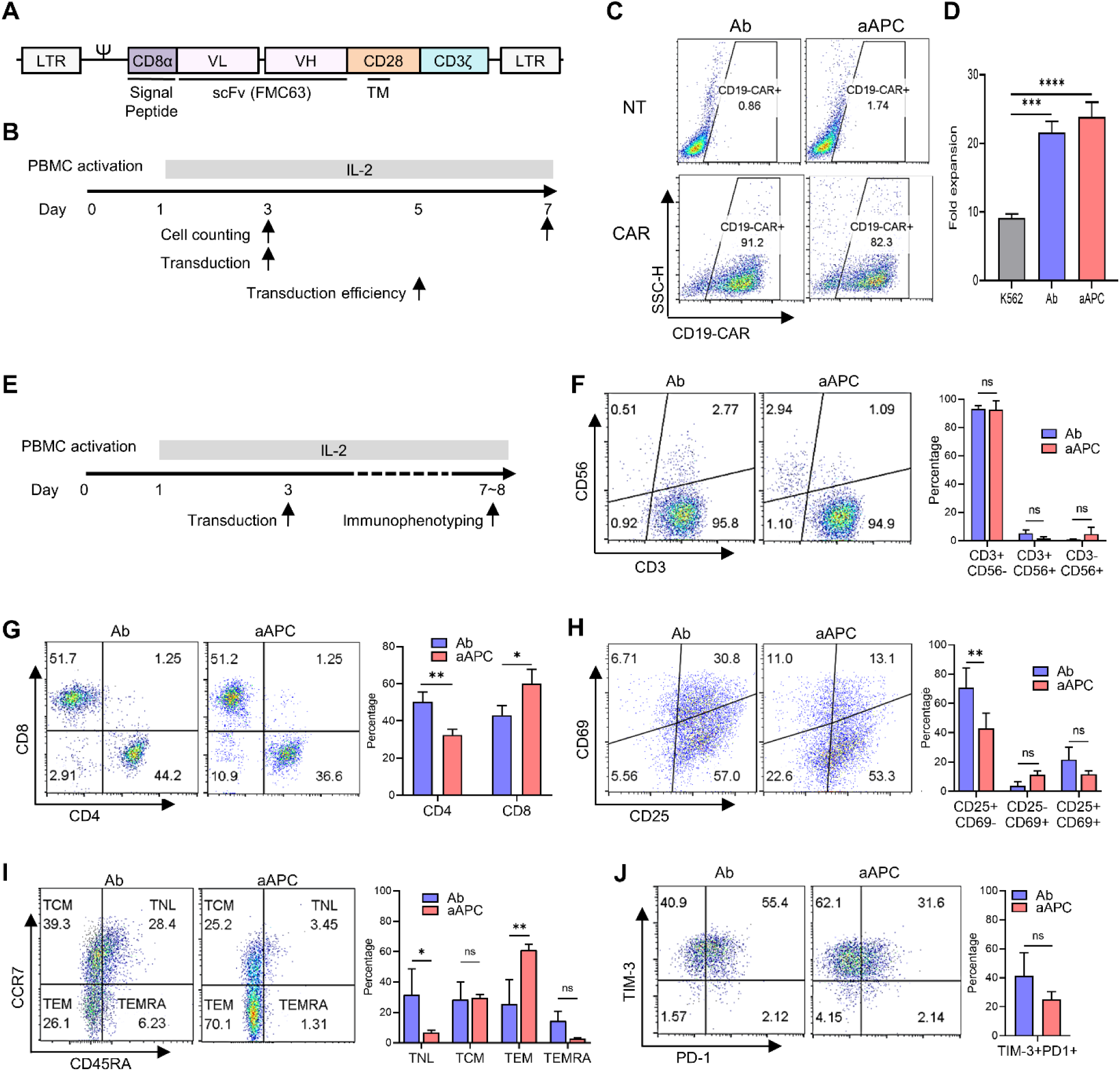
Design of CD19-CAR structure and effect of aAPC and Ab activation methods on the expansion of CD19-CAR-positive cells and evaluation of their immunophenotype. (A) Schematics of the anti-CD19 CAR construct with single-chain variable fragment (scFv) component (anti-CD19 antibody FMC63), CD28 co-stimulatory signaling domain, and CD3ζ. (B) Experimental timeline. (C) Cell surface CAR expression on transduced cells with CD19-CAR construct was determined by flow cytometry and comparison of transduction efficiency of CAR positive cells activated by aAPC and antibody (Ab) methods. (D) Fold change of activated CD19-CAR positive cells with aAPC cell, anti-CD3/CD28 antibodies, and K562 cell line. Cell expansion was assessed at day 3 to day 7. (E) Experimental timeline of T cell activation, transduction, cell expansion, and staining. (F) Staining of cells with anti-CD56 and anti-CD3 to determine T and NK cell populations. Similar to the Ab activated method, aAPC cells can expand CAR T cells (CD3^+^ CD56^−^) and have no significant changes in the expansion of the NKT population (CD3^+^CD56^+^). (G) Analysis of T cells showed that CD8^+^ CAR T cells were more expanded in the aAPC activated method and CD4^+^ CAR T cells in the Ab-activation method. (H) Flow cytometry analysis of CD25 and CD69 activation markers of CAR T cells in aAPC and Ab activated method. The data showed no difference between the expressions of CD25^+^CD69^+^ dual activation markers and the expression of CD25^+^CD69^−^ marker increased in Ab activated method. (I) Representative flow cytometry and quantification data for memory phenotype of CAR T cells on day 8. In the aAPC activated method, the differentiation of CAR T cells to TEM cells increased, and in the Ab activated method, the differentiation of CAR T cells to TNL cells increased. (J) The expression of PD-1 and TIM-3 exhaustion markers of CAR T cell with different activation methods showed that the exhaustion markers were not significant in both groups. TNL: naïve-like T cell; TCM: central memory T cell; TEM: effector memory T cell; TEMRA: CD45RA^+^ effector memory T cell. *n* = 3. * P < 0.05; ** P < 0.01; *** P < 0.001; **** P < 0.0001.

### Effect of aAPCs-mediated activation on the immunophenotype of CAR T cells

To evaluate the expansion of CD3^+^ T cell and NK cell population on day 3, activated T cells were transduced with CD19-CAR bearing retroviral, and CD3 and CD56 antibodies were used for cell staining on day 7. The data indicated that the aAPC activated group induced the same expansion in CD3^+^ T cells as the Ab activated group, and there was no significant change in expansion in the NKT population (CD3^+^ CD56^+^) in the two groups (Fig. 4E and F). Similar to what was observed with untransduced T cells, aAPCs expanded cytotoxic CD8^+^ CAR T cells more than Ab group (Fig. 4G). We did not observe any difference in CD25^+^CD69^+^ CAR T cell population between aAPC and Ab groups while the percentage of CD25^+^CD69^−^ CAR T cell was lower in aAPC group (Fig. 4H). Although this result was consistent with the previous experiment with untransduced T cells (Fig. 3E), the magnitude of difference in CD25^+^CD69^−^ sub-population between the two activation methods was lower in the case of CAR transduced T cells (1.6 fold) compared to untransduced T cells (>5 fold). As presented in Fig. 4I, aAPC-mediated activation method resulted in increased TEM (CD45RA^−^CCR7^−^) and reduced TNL (T naïve like; CD45RA^+^CCR7^+^) percentages among CAR T cell population compared with the Ab-mediated activation method. No statistically significant difference was observed in TCM (CD45RA^−^CCR7^+^) and TEMRA (CD45^+^CCR7^−^) sub-populations. Moreover, the prevalence of PD-1 and TIM-3 expressing CAR T cells was not different between the two experimental groups (Fig. 4J).

### Assessment of activation and exhaustion characteristics of CAR T cells and the ability of antitumor activity

In order to evaluate the effect of aAPC-activation method on the CAR-mediated activation of CD19-CAR T cells, we co-cultured CAR T cells or untransduced T cells with CD19^+^ target cells (Raji cells) at a ratio of 1:1 (Fig. 5A). After 24 hours, we observed an extremely high IFN-γ level in the supernatant of co-cultured CAR T cells compared to untransduced T cells in aAPC and Ab groups. Also, CAR T cells activated with aAPC produced significantly higher IFN-γ compared to CAR T cells activated with ab (Fig. 5B). Also, CAR T cells activated with both aAPC and Ab showed increased expression of CD25 and CD69 activation markers compared with untransduced T cells (Fig. 5C). To measure CAR T cell exhaustion, a longer co-culture (16 days) with a higher number of target cells (effector to target ratio of 1:5) was used (Fig. 5A). The results showed that in both aAPC and Ab CAR T cell groups higher percentage of cells co-expressed PD-1 and TIM-3 compared with the untransduced T cells, however, there was no difference between aAPC and Ab activation conditions (Fig. 5D). Also, the results showed that the expression of TIM-3 in both groups of CAR T cells was higher than non-transduced T cells, although PD-1 expression was not different in the CAR T cells and non-transformed T cells (Fig. S1).

**Figure 5.**
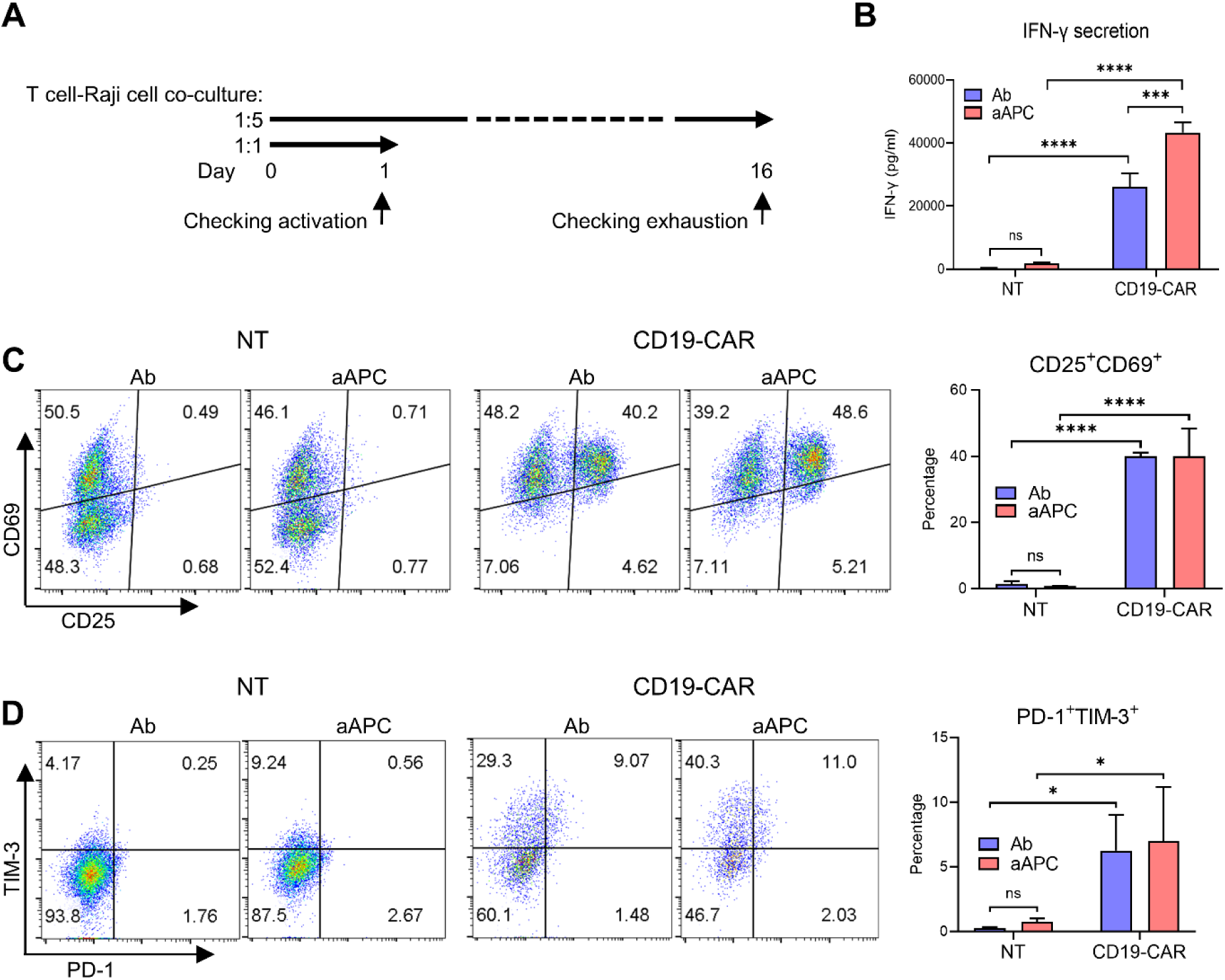
Antitumor effects and expression of activation and exhaustion markers in co-culture with target cells. (A) Experimental timeline of co-culture of CAR T cells with Raji cells. (B) Co-culture of CAR T cells with Raji cells at a ratio of 1:1 and collection of supernatants to evaluate IFN-γ release assay after 24 h. The data showed that aAPC activated CAR T cells significantly produced IFN-γ compared with antibody (Ab) activated CAR T cells. (C) CAR T cells and Raji cells were co-cultured at a ratio of 1:1 and after an overnight expression of CD25 and CD69 were measured by flow cytometry. The data showed that aAPC activated method similar to Ab activated method can express activation markers. (D) CAR T cells and Raji cells were co-cultured at a ratio of 1:5 (E:T). After 16 days, CAR T cells activated with aAPC cells express exhaustion markers PD-1 and TIM-3 upon long-term exposure to Raji cells, similar to Ab activated method. *n* = 3. * P < 0.05; *** P < 0.001; **** P < 0.0001.

### Cytotoxicity of CAR T cells generated by aAPC-mediated activation

To investigate the cytotoxic function of aAPC CAR T cells, they co-cultured with FFluc^+^ Raji cells at effector to target (E:T) ratios of 10:1, 5:1: 2.5:1, 1:1 and 0:1 in 96-well flat-bottom plates. Co-culture of untransduced T cells with tumor cells and tumor cells without effector cells were performed as a control. After 6 hours incubation cytotoxicity was measured in a luciferase-based assay. The results showed indicated that E:T-dependent cytotoxicity for both aAPC and Ab-drived CD19-CAR T cells compared with untransduced T cells without any significant difference between the cytotoxic activity of CAR T cells generated by aAPC and Ab (Fig. 6A and B).

**Figure 6.**
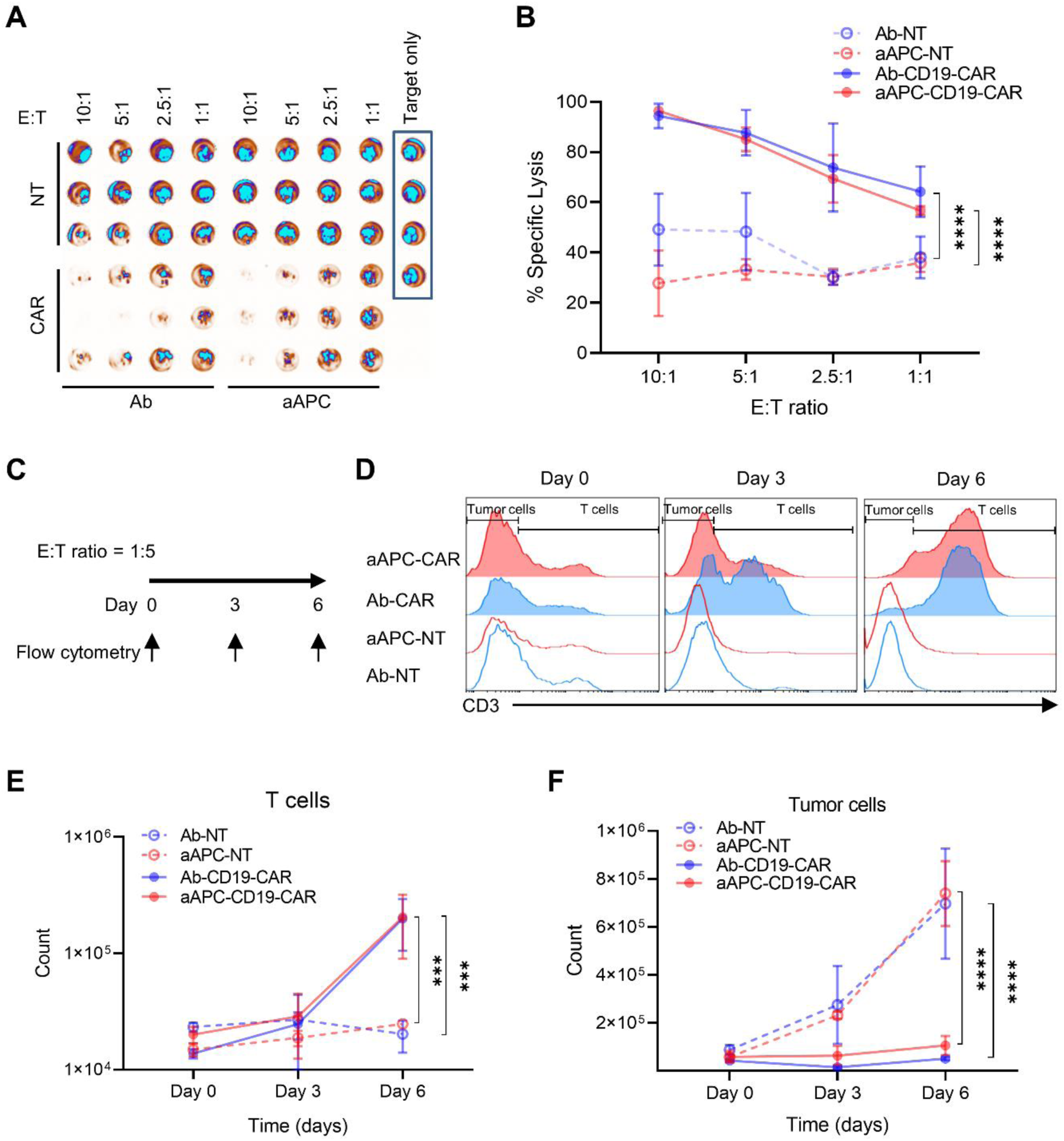
In vitro anti-tumor response of CD19-CAR T cells. (A and B) Specific tumor cell lysis by CD19-CAR T cells. CD19-CAR T cells (CAR) and untransduced T cells (NT) were co-cultured with ffluc-expressing Raji cells at different E:T ratios and after 6 hours of incubation cell lysis was measured in a luciferase-based assay. The results indicated that specific lysis occurred with increasing numbers of CD19-CAR T cells, and aAPC activated CAR T cells had similar specific lysis activity to antibody (Ab) activated CAR T cells. (A) Experimental timeline of co-culture of CAR T cells with Raji cells for specific lysis and cytotoxicity. (D) Representative flow cytometry histogram showing the cytotoxicity of CD19-CAR T cells at an E:T ratio of 1:5 for 6 days. Cells were harvested every 3 days for staining with anti-CD3 and the percentage of cell viability was measured with 7AAD. CAR T-mediated killing assays at 6 days after co-culture showed that the majority of Raji cells were eliminated. (E and F) Number of T cells and tumor cells during co-culture. (E) Proliferation of CD19-CAR T cells from panel D cultures occurred significantly higher than NT T cells. (F) The number of eliminated Raji cells was significantly increased in co-culture with CD19-CAR T cells from panel D compared to co-culture with NT cells. The behavior of aAPC activated CAR T cells in co-culture with tumor cells was comparable to that of Ab activated CAR T cells. *n* = 3. *** P < 0.001; **** P < 0.0001

To further investigate the anti-tumor capacity of CAR T cells, a 6-day co-culture was performed with Raji cells at a higher ratio of target cells with an E:T of 1:5 (Fig. 6C). The T cells and Raji calls were detected on days 0, 3 and 6 by CD3 staining (Fig. 6D). Absolut quantification of the cell numbers using counting beads showed that both AB and aAPC-drived CAR T cells expanded in the course of co-culture while untransduced T cells did not proliferate (Fig. 6E). Additionally, the number of Raji cells were decreased in both AB and aAPC-drived CAR T cell groups, unlike the untransduced groups which showed Raji cell growth (Fig. 6F). In summary, both CAR T cells activated with aAPC line and Ab were able to proliferate upon exposure to tumor cells and control the target cells with statistically indifferent rates.

## Discussion

The importance of CAR T cell therapy compared to other treatments such as chemotherapy and conventional immunotherapies is that it shows better results in patients who have had multiple relapses (21, 22). In this treatment method, multiple infusions are not needed, and despite this, the treatment period is shorter and recovery is faster. So far, CAR T cell therapy for some blood cancers has been able to overcome the limitations of other conventional treatments. Among current therapies, checkpoint blockade has shown better results, especially in solid tumors. However, combination therapy of CAR T cells with drugs such as checkpoint blockade has been proposed and may provide good results (21–24).

In order to improve the activation and expansion of T cells for high-scale cultures, the use of aAPCs is a suitable and cost-effective approach (11–13). After dendritic cells (DC) were identified in the 1970s (25), one of the first strategies was to use them as natural APCs, which sense antigens and act as mediators for other immune cells, especially T cells (26). DCs stimulate TCR and co-stimulatory signals on T cells through MHC expression molecules and are a strong stimulus for T cell activation (27, 28). Many researchers have generated different types of engineered DCs to better expand T cells, but limitations of its use included inconsistency of antitumor responses among donors, reduced functionality and longevity of memory cells generated by this method (29–31). Immunologists then successfully expanded human T cells with aAPCs instead of natural APCs. The aAPC-based system is one of the strategies used to activate and expand T cells to generate CAR T cells for adoptive cell transfer. The use of K562 aAPC for T cell activation has many advantages besides being cost-effective, not expressing MHC and inhibitory molecules, including additional stimulatory molecules that interact more with T cells (10, 11).

Anna K. Thomas and colleagues generated aAPCs that expressed CD3/CD28, which more rapidly induced T cells to enter the cell cycle and after 10 days of culture, expanded T cells fourfold more than CD3/CD28-coated beads (15). By co-culturing aAPCs with PBMCs in different ratios, a cluster of aAPC and T cells was formed.

In our study, we engineered a K562 aAPC by constructing a membrane-bound OKT3 retroviral plasmid encoding OKT3-GFP. By co-culturing aAPCs with PBMCs in different ratios, clusters of aAPC and T cells were formed. We observed that large clusters were formed in the ratio of 1:1, but at a 1:0.5 ratio the size of clusters formed was similar to anti-CD3/CD28 standard T cell activation method (Fig. 2C). Since one of the important considerations in using aAPC for T cell stimulation is the duration of stimulation and persistent signaling causes cell exhaustion (32), we needed to investigate the disappearance of aAPCs in co-culture. In a previous study, the disappearance of irradiated aAPC from the co-culture was measured and adjusted at a ratio of 2:1, and the number of aAPC was determined by total cell count and flow cytometry. The number of irradiated aAPCs in the co-culture started to decrease on the first day and reached zero on the third day. In our study we examined the disappearance of mitomycin C-treated aAPC in co-culture from day 0 to day 2 by total cell count and flow cytometry. We observed that on day 2, at a ratio of 1:1, some aAPCs were still alive, but at ratios of 1:0.5 and 1:0.25, aAPCs are almost eliminated by T cells. Due to the similarity of the size of the clusters in the ratio of 1:0.5 with the Ab activated method and the disappearance aAPC on day 2, we chose this ratio for the next experiments.

The results shown in Fig. 3C and Fig. 4F that activation of T cells with aAPC expands CD3^+^ T cells similar to Ab activation method, but in aAPC activation method compared to Ab activation method, CD8^+^ T cells expand significantly more than CD4^+^ T cells (Fig. 3D and 4G). When CD19-CAR T cells produced by these methods were examined, these results were confirmed similar to untransduced T cells. Our finding confirmed a previous result by Butler et al (33), that aAPC/mOKT3 promoted the expansion of CD8^+^ T cells better than CD4^+^ T cells. Bishwas Shrestha and colleagues replicated similar results while generating aAPCs expressing anti-CD3, anti-CD28, and in combination with CD137L (34), whereas our generated aAPCs express only anti-CD3, causing this increase in CD8^+^ T cell expansion.

When we evaluated T cell activation markers on day 7 without co-culture with tumor cells, there was no difference between the expression level of CD25^+^CD69^+^ double positive and double negative markers, but the expression of CD25^+^CD69^−^ is increased in Ab activation method rather than aAPC activation method (Fig. 3E). In a previous study, we found that after stimulation of T cells with anti-CD28, the expression level of CD25 was increased (35) and another study indicated that CD25 gene expression in T cells is rapidly induced upon TCR/CD28 co-stimulation by activation of NF-κB, NFAT, AP-1, and cAMP response element-binding protein/activating transcription factor (36). It is clear that our aAPCs express only anti-CD3 and lack the expression of CD28, and therefore the increase in CD25 expression was not the same as in the Ab activation group. It is important that the expression of activation markers increases after the exposure of T cell and CAR T cell to the tumor cell. As shown in figure 5C, the expression of these markers was significantly increased in CAR T cells activated with aAPC and Ab activated methods after exposure to tumor cells in a similar manner compared to the untransduced T cells.

To evaluate the effect of different activation methods on T cell memory phenotype, we analyzed untransduced T cells on day 8. The results indicate that aAPC activation method increased the population of TEM cells, but Ab activation method increased TNL cell population. It seems that exposure of T cells with aAPCs in the aAPC activation method has caused the differentiation of T cells in this group more towards the memory effector. There was no significant difference in the cell population of TCM (T central memory) and TEMRA (Terminally differentiated effector memory T) in the two activation methods. In the study they used CD3/28/137L K562 aAPCs did not observe a significant difference in CD8 TCM and CD8 TEM between CD3/28/137L and aAPC beads, but they showed that activation of T cells with aAPCs resulted in a higher percentage of memory CD8 T cells (34). Also, Kawalekar et al observed similar results where CAR T cells with 41BB/CD137L endodomain increased their oxidative phosphorylation characteristic of CD8 memory T cells (37). In contrast to these two groups, activation with our aAPCs showed an increase in TEM cells in the total population of non-transduced T cells and CAR T cells. On day 8, we evaluated the expression of PD-1 and TIM-3 exhaustion markers on T cells in two activation methods and the findings showed that the exhaustion markers of the aAPC method are lower, but there is no significant difference in the expression level of the exhaustion markers with the Ab activation method. This finding showed that aAPCs do not have a great effect on T cell exhaustion like the standard activation method. In a previous study using K562 CD3/28/137L aAPCs, intrinsic glycolysis (ECAR) and oxidative phosphorylation mitochondrial index (OCAR) of CD8^+^ T cells were measured to assess T cell exhaustion. They showed that CD8^+^ T cells activated with aAPC CD3/28/137L K562 cells have a higher glycolytic phenotype, along with increased mitochondrial oxidation (34). Compared to this study, our aAPC only expressed anti-CD3 and also the type of cellular exhaustion measurement is different from our method, and this group only investigated cellular exhaustion on CD8^+^ T cells, but we measured it on the whole cell population. The memory phenotype and exhaustion marker of CD19-CAR T cells produced by aAPC and Ab activation methods confirmed the results of untransduced T cells.

To further assess antigen-mediated T cell activation, we exposed CD19-CAR T cells with Raji target cells at a 1:1 ratio. After overnight stimulation, measured upregulation of the surface marker CD25/CD69. The finding showed that the CAR T cell produced by aAPC method showed a strong upregulation of CD25/CD69 when co-cultured with target cells as well as the Ab activation method. The results indicate that CAR T cells are activated by aAPC when exposed to antigen-presenting cells, expressing specific activation markers as in the standard method. The results of the IFN-γ release assay of CAR T cells produced by both methods compared to untransduced T cells showed that CAR T cells were able to secrete a high level of IFN-γ after co-culture with the target cells (Fig. 5B). Interestingly, the CAR T cells produced by the aAPC method significantly produced IFN-γ compared to the Ab activation method, which is consistent with the result presented in the Fig. 4I, which showed that aAPC activation method induced T cells to further differentiate into effector memory (TEM) phenotype. Also when co-cultured aAPC activated CAR T cells with Raji cells at a 1:5 E:T ratio and culture continued for 16 days, no difference with Ab activated CAR T cells in the expression level of PD-1 and TIM-3 exhaustion markers. In a study in which aAPC produced anti-CD3 and anti-CD28 in combination with CD137L, compared to T cells stimulated with anti-CD3/CD28 beads, the aAPC-expanded CD8 T cell subpopulation expressed fewer markers of exhaustion (34), instead, we did not measure exhaustion markers in T cell sub-populations and only in the total T cell population.

In addition, we co-cultured CAR T cells with ffluc-expressing Raji cells at various E:T ratios and measured antitumor immune responses in vitro in a luciferase-based assay. We showed that CAR T cells activated with aAPC had ∼95% specific lysis potency, and no significant lysis was observed for untransduced T cells. Also, the results show that CAR T cells activated with aAPC method have the same potency as CAR T cells activated by Ab (Fig. 6A and B). Also, we evaluated the anti-tumor capacity of CAR T cells by co-culture with Raji target cell at a higher E:T ratio (1:5) at 3-time points. As it is clear in the results, the aAPC activated CAR T cells eliminated the target cells in a period of 6 days, and similar to the standard method, the activated aAPC CAR T cells were equivalent to the standard method in terms of killing potential. We also counted the number of T cells and tumor cells by adding counting beads to the cell mixture of co-culture and found that aAPC and Ab activated CAR T cell subsets were able to proliferate when exposed to tumor cells, unlike untransduced T cells. The result showed that the proliferation of tumor cells co-cultured with untransduced T cells increased, but in the presence of both groups of CAR T cells, the tumor cells were killed and proliferation did not occur (Fig. 6D-F). Andrea Schmidts and colleagues that used CD3^+^CD28^+^K562 aAPC as a T cell activator, checked in vitro potency of aAPC activated CD19-CAR T cells on Nalm6 and Jeko-1 target cells by luciferase-based assay and showed aAPC activated CD19-CAR T cells similar to anti-CD3/CD28 bead activated CD19-CAR T cells have potency in killing tumor cells (14). Compared to this study, in addition to the luciferase-based method, we used long-term co-culture and a high ratio of target cells to measure CAR T cell potency as well as CAR T cell and target cell proliferation. In a study, K562 aAPCs were generated that expressed anti-CD3/CD28 and, in addition, disrupted endogenous low-density lipoprotein receptor (LDLR) gene expression in these aAPCs to prevent unwanted lentiviral transduction. They hypothesized that K562 cells are sensitive to transduction by lentiviral vectors due to constitutive expression of LDLR, and unwanted aAPC transduction can reduce T cell transduction (14). In the current study, we used aAPC for the activation of T cell and after removing them by T cells, on day 3, we transduced T cells and observed a high rate of cell transduction (80-90%) as in the standard method of Ab activation. Therefore, our generated aAPC can be used as a candidate for activation and preparation of T cells for transduction as in the standard method without reducing the transduction rate.

The safety of K562 cell was previously demonstrated in a phase 2 AML study in which autologous leukemia cells were mixed with GM-CSF-secreting K562 cells and primed lymphocytes re-infused with stem cell transplantation (18). In another study, patients with CML received a series of intradermal vaccines containing irradiated K562/GM-CSF cells (19). Based on these trial studies that evaluated the safety of the irradiated-K562, it is hoped that engineered K562 cells can also be used for T cell activation with cost-effective production and further expansion of CD8^+^ T cells.

## Conclusion

In summary, our findings suggest that the engineered OKT3 aAPC, as simpler and more cost-effective, has the potential to expand large numbers of CAR T cells for immunotherapy of cancer patients. It was also shown that after the second day, we have no residual aAPCs in the culture, so it does not interfere with the transduction of T cells, and the rate of transduction is comparable to the Ab activation method. It was also shown that this aAPC does not change the phenotype and exhaustion level of the CAR T cells and increases the proliferation of CD8^+^ killer cells compared to the standard method. CAR T cells expanded by this method after exposure to target cells show significant expression of activation markers, and the potential of these cells in specific lysis and cytotoxicity in co-culture with a high proportion of target cells is competitive with the standard method. This aAPC cell has the potential to activate and expand T cells and is also proposed as a cost-effective way to generate CAR T cells for cancer studies. Overall, our results show that engineered aAPC cells similar to the standard Ab activation method can activate T cells and even further expand CD8+ T cells, so this method is introduced as a cost-effective way to generate CAR T cells for cancer immunotherapy.

## Author Contributions

Conceptualization, A.S. and M.B; methodology, A.S., M.A., and M.B.; data analysis, A.S. and M.B.; writing, review and editing the manuscript, A.S., A.A., K.Y., H.S., D.S., B.B., and M.B.; supervision and funding acquisition, B.B. and M.B.

## Institutional Review Board Statement

This study was approved by the Academic Research Ethics Committee of Tabriz University of Medical Sciences (code of ethics, IR. TBZMED. REC. 1400.951). Informed consent was obtained from all blood donors involved in the study.

## Acknowledgments

We thank all staff and technicians at Royan Institute for Stem Cell Biology and Technology for their technical assistance especially Gene Therapy and CAR T cell team members. This research was supported by Tabriz University of Medical Sciences (Grant code. 68848).

## References

1. Rosenberg SA, Restifo NP. Adoptive cell transfer as personalized immunotherapy for human cancer. Science. 2015;348(6230):62–8.

2. Redeker A, Arens R. Improving adoptive T cell therapy: the particular role of T cell costimulation, cytokines, and post-transfer vaccination. Frontiers in immunology. 2016;7:345.

3. Maus MV, Fraietta JA, Levine BL, Kalos M, Zhao Y, June CH. Adoptive immunotherapy for cancer or viruses. Annual review of immunology. 2014;32:189–225.

4. Mullard A. FDA approves first CAR T therapy. Nature reviews Drug discovery. 2017;16(10):669.

5. Morgan RA, Dudley ME, Wunderlich JR, Hughes MS, Yang JC, Sherry RM, et al. Cancer regression in patients after transfer of genetically engineered lymphocytes. Science. 2006;314(5796):126–9.

6. Noonan KA, Huff CA, Davis J, Lemas MV, Fiorino S, Bitzan J, et al. Adoptive transfer of activated marrow-infiltrating lymphocytes induces measurable antitumor immunity in the bone marrow in multiple myeloma. Science translational medicine. 2015;7(288):288ra78.

7. Xu D, Jin G, Chai D, Zhou X, Gu W, Chong Y, et al. The development of CAR design for tumor CAR-T cell therapy. Oncotarget. 2018;9(17):13991.

8. June CH, Sadelain M. Chimeric antigen receptor therapy. New England Journal of Medicine. 2018;379(1):64–73.

9. Sterner RC, Sterner RM. CAR-T cell therapy: current limitations and potential strategies. Blood cancer journal. 2021;11(4):69.

10. Safarzadeh Kozani P, Safarzadeh Kozani P, Rahbarizadeh F. CAR-T cell therapy in T-cell malignancies: Is success a low-hanging fruit? Stem cell research & therapy. 2021;12(1):1–17.

11. Iyer RK, Bowles PA, Kim H, Dulgar-Tulloch A. Industrializing autologous adoptive immunotherapies: manufacturing advances and challenges. Frontiers in medicine. 2018;5:150.

12. McCarron A, Donnelley M, McIntyre C, Parsons D. Challenges of up-scaling lentivirus production and processing. Journal of biotechnology. 2016;240:23–30.

13. Merten O-W, Hebben M, Bovolenta C. Production of lentiviral vectors. Molecular Therapy-Methods & Clinical Development. 2016;3:16017.

14. Schmidts A, Marsh LC, Srivastava AA, Bouffard AA, Boroughs AC, Scarfò I, et al. Cell-based artificial APC resistant to lentiviral transduction for efficient generation of CAR-T cells from various cell sources. Journal for immunotherapy of cancer. 2020;8(2).

15. Thomas AK, Maus MV, Shalaby WS, June CH, Riley JL. A cell-based artificial antigen-presenting cell coated with anti-CD3 and CD28 antibodies enables rapid expansion and long-term growth of CD4 T lymphocytes. Clinical immunology (Orlando, Fla). 2002;105(3):259–72.

16. Maus MV, Thomas AK, Leonard DG, Allman D, Addya K, Schlienger K, et al. Ex vivo expansion of polyclonal and antigen-specific cytotoxic T lymphocytes by artificial APCs expressing ligands for the T-cell receptor, CD28 and 4-1BB. Nature biotechnology. 2002;20(2):143–8.

17. Prakken B, Wauben M, Genini D, Samodal R, Barnett J, Mendivil A, et al. Artificial antigen-presenting cells as a tool to exploit the immunesynapse’. Nature medicine. 2000;6(12):1406–10.

18. Borrello IM, Levitsky HI, Stock W, Sher D, Qin L, DeAngelo DJ, et al. Granulocyte-macrophage colony-stimulating factor (GM-CSF)–secreting cellular immunotherapy in combination with autologous stem cell transplantation (ASCT) as postremission therapy for acute myeloid leukemia (AML). Blood, The Journal of the American Society of Hematology. 2009;114(9):1736–45.

19. Qin L, Smith B, Tsai H, Yaghi N, Neela P, Moake M, et al. Induction of high-titer IgG antibodies against multiple leukemia-associated antigens in CML patients with clinical responses to K562/GVAX immunotherapy. Blood cancer journal. 2013;3(9):e145-e.

20. Yekehfallah V, Pahlavanneshan S, Sayadmanesh A, Momtahan Z, Ma B, Basiri M. Generation and Functional Characterization of PLAP CAR-T Cells against Cervical Cancer Cells. Biomolecules. 2022;12(9):1296.

21. Want MY, Bashir Z, Najar RA. T Cell Based Immunotherapy for Cancer: Approaches and Strategies. Vaccines. 2023;11(4):835.

22. Sheykhhasan M, Manoochehri H, Dama P. Use of CAR T-cell for acute lymphoblastic leukemia (ALL) treatment: a review study. Cancer gene therapy. 2022;29(8-9):1080–96.

23. Houot R, Schultz LM, Marabelle A, Kohrt H. T-cell–based immunotherapy: adoptive cell transfer and checkpoint inhibition. Cancer immunology research. 2015;3(10):1115–22.

24. Ramos CA, Heslop HE, Brenner MK. CAR-T cell therapy for lymphoma. Annual review of medicine. 2016;67:165–83.

25. Steinman RM, Cohn ZA. Identification of a novel cell type in peripheral lymphoid organs of mice: I. Morphology, quantitation, tissue distribution. The Journal of experimental medicine. 1973;137(5):1142–62.

26. Steinman RM. The dendritic cell system and its role in immunogenicity. Annual review of immunology. 1991;9:271–96.

27. Inaba K, Metlay JP, Crowley MT, Steinman RM. Dendritic cells pulsed with protein antigens in vitro can prime antigen-specific, MHC-restricted T cells in situ. J Exp Med. 1990;172(2):631–40.

28. Ye K, Li F, Wang R, Cen T, Liu S, Zhao Z, et al. An armed oncolytic virus enhances the efficacy of tumor-infiltrating lymphocyte therapy by converting tumors to artificial antigen-presenting cells in situ. Molecular Therapy. 2022;30(12):3658–76.

29. Ratta M, Fagnoni F, Curti A, Vescovini R, Sansoni P, Oliviero B, et al. Dendritic cells are functionally defective in multiple myeloma: the role of interleukin-6. Blood, The Journal of the American Society of Hematology. 2002;100(1):230–7.

30. Satthaporn S, Robins A, Vassanasiri W, El-Sheemy M, Jibril JA, Clark D, et al. Dendritic cells are dysfunctional in patients with operable breast cancer. Cancer Immunology, Immunotherapy. 2004;53:510–8.

31. Neal LR, Bailey SR, Wyatt MM, Bowers JS, Majchrzak K, Nelson MH, et al. The Basics of Artificial Antigen Presenting Cells in T Cell-Based Cancer Immunotherapies. Journal of immunology research and therapy. 2017;2(1):68–79.

32. Wherry EJ. T cell exhaustion. Nature immunology. 2011;12(6):492–9.

33. Butler MO, Imataki O, Yamashita Y, Tanaka M, Ansén S, Berezovskaya A, et al. Ex vivo expansion of human CD8+ T cells using autologous CD4+ T cell help. PloS one. 2012;7(1):e30229.

34. Shrestha B, Zhang Y, Yu B, Li G, Boucher JC, Beatty NJ, et al. Generation of antitumor T cells for adoptive cell therapy with artificial antigen presenting cells. Journal of Immunotherapy (Hagerstown, Md: 1997). 2020;43(3):79.

35. Vidan MT, Fernandez-Gutierrez B, Hernandez-Garcia C, Serra J, Ribera J, Perez-Blas M, et al. Functional integrity of the CD28 co-stimulatory pathway in T lymphocytes from elderly subjects. Age and ageing. 1999;28(2):221–7.

36. Beyersdorf N, Kerkau T, Hünig T. CD28 co-stimulation in T-cell homeostasis: a recent perspective. ImmunoTargets and therapy. 2015:111–22.

37. Kawalekar OU, O’Connor RS, Fraietta JA, Guo L, McGettigan SE, Posey AD, Jr., et al. Distinct Signaling of Coreceptors Regulates Specific Metabolism Pathways and Impacts Memory Development in CAR T Cells. Immunity. 2016;44(2):380–90.

