## Supplementary Figure for "Characterization of CAR T Cells Manufactured using Genetically Engineered Artificial Antigen Presenting Cells"

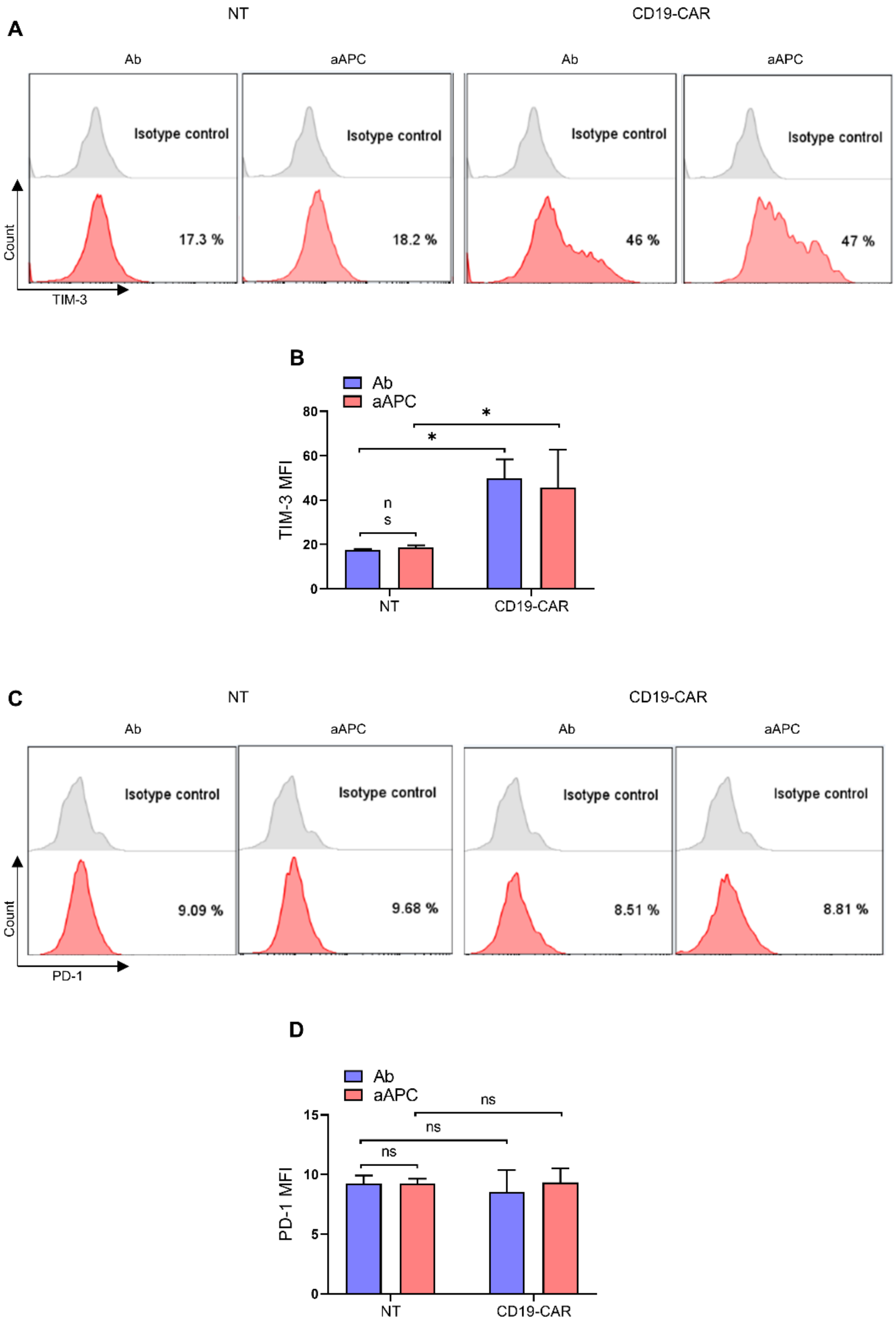


**Figure S1. Analysis of the PD-1 and TIM-3 expression markers separately in co-culture with target cells.** The analysis of PD-1 and TIM-3 expression markers separately showed that TIM-3 marker was higher in CAR T cell groups than not-transduced T cells (A and B), but the difference of PD-1 marker expression in CAR T cell groups and non-transduced T cells are not significant (C and D).
